# High-resolution fecal pharmacokinetic modeling in mice with orally administered antibiotics

**DOI:** 10.1101/2023.05.05.539495

**Authors:** Rie Maskawa, Lena Takayasu, Hideki Takayasu, Keiji Watanabe, Shusuke Takemine, Takashi Kakimoto, Kozue Takeshita, Seiko Narushima, Wataru Suda, Misako Takayasu

## Abstract

High-resolution fecal pharmacokinetics are crucial for optimizing therapeutic design and evaluating gastrointestinal motility. However, empirical studies with detailed time series data remain limited. This study aims to characterize fecal pharmacokinetics through high-frequency sampling and parallelized fecal concentration quantification, establishing a simple pharmacokinetics model with physiologically interpretable parameters. We quantified vancomycin concentrations in fecal samples collected at a minimum interval of 4 hours from C57BL/6J mice following a single oral administration of either a low (1 mg/mL) or high (20 mg/mL) dose. Fecal concentrations gradually increased and exhibited an exponential decay, leading to the development of a compartmental model with an absorption phase. This simple model accurately fit the experimental data and provided physiological explanations for intra- and inter-individual pharmacokinetics variability. The results suggest that inter-individual differences in pharmacokinetics are attributable to fecal elimination capacity, which may be influenced by drug dosage via changes in gastrointestinal motility. Since the model predicts antibiotic concentrations within the gastrointestinal tract, it can be applied to fundamental studies investigating the effects of antibiotics on the gut microbiome and gastrointestinal motility.

## Introduction

Fecal pharmacokinetics plays a crucial role in optimizing therapeutic strategies and assessing gastrointestinal motility. However, empirical studies using high-resolution time series data remain limited, and detailed information on the temporal profiles and inter-individual variability of fecal drug concentrations has not been fully elucidated. In particular, accurate assessment of fecal drug concentrations after oral administration is essential for understanding the gastrointestinal pharmacokinetics and elimination mechanisms of drugs. To date, pharmacokinetic parameters have been derived from time series of fecal drug concentrations using non-parametric methods^1,2^.

Vancomycin (VCM) is a glycopeptide antibiotic used primarily to treat Gram-positive bacterial infections^3^. When administered orally, systemic absorption is minimal, resulting in high fecal concentrations^4^. In human patients with *Clostridium difficile* infection, fecal VCM concentrations and stool frequency have been measured, and the relationship between dosage and fecal drug concentrations has been discussed^4,5^. Despite its clinical importance, quantitative studies of its fecal pharmacokinetics with high temporal resolution have not been widely performed. In addition, conventional nonparametric analyses often fail to explain the underlying kinetic mechanisms that account for the variability in drug transport and excretion^6–10^. Therefore, to deepen our understanding of drug behavior in the gastrointestinal tract, it is necessary to develop pharmacokinetic models that are physiologically interpretable.

In this study, we characterized the fecal pharmacokinetics of orally administered VCM in C57BL/6J mice using high-frequency fecal sampling and parallel quantification of fecal drug concentrations. We aimed to establish a physiologically interpretable framework that accurately describes drug transport dynamics by constructing a minimal pharmacokinetic model that captures the robustly observed data characteristics.

## Results

### Fecal Pharmacokinetics of Vancomycin

To evaluate the detailed temporal changes in fecal VCM concentrations, we performed high-frequency fecal sampling and parallel quantification of antibiotic concentrations (Figure 1). At the initial time point (day 0, 12:00), C57BL/6J mice in the low-dose group (1 mg/mL, L1–L3) and high-dose group (20 mg/mL, H1–H3) received an oral administration of 0.5 mL of VCM. Fecal samples were collected at a minimum interval of 4 hours. After purification using solid-phase extraction columns, VCM concentrations in feces were calculated by normalizing the mass measured by LC-MS/MS to the fecal mass (see Methods for details).

**Figure 1.**
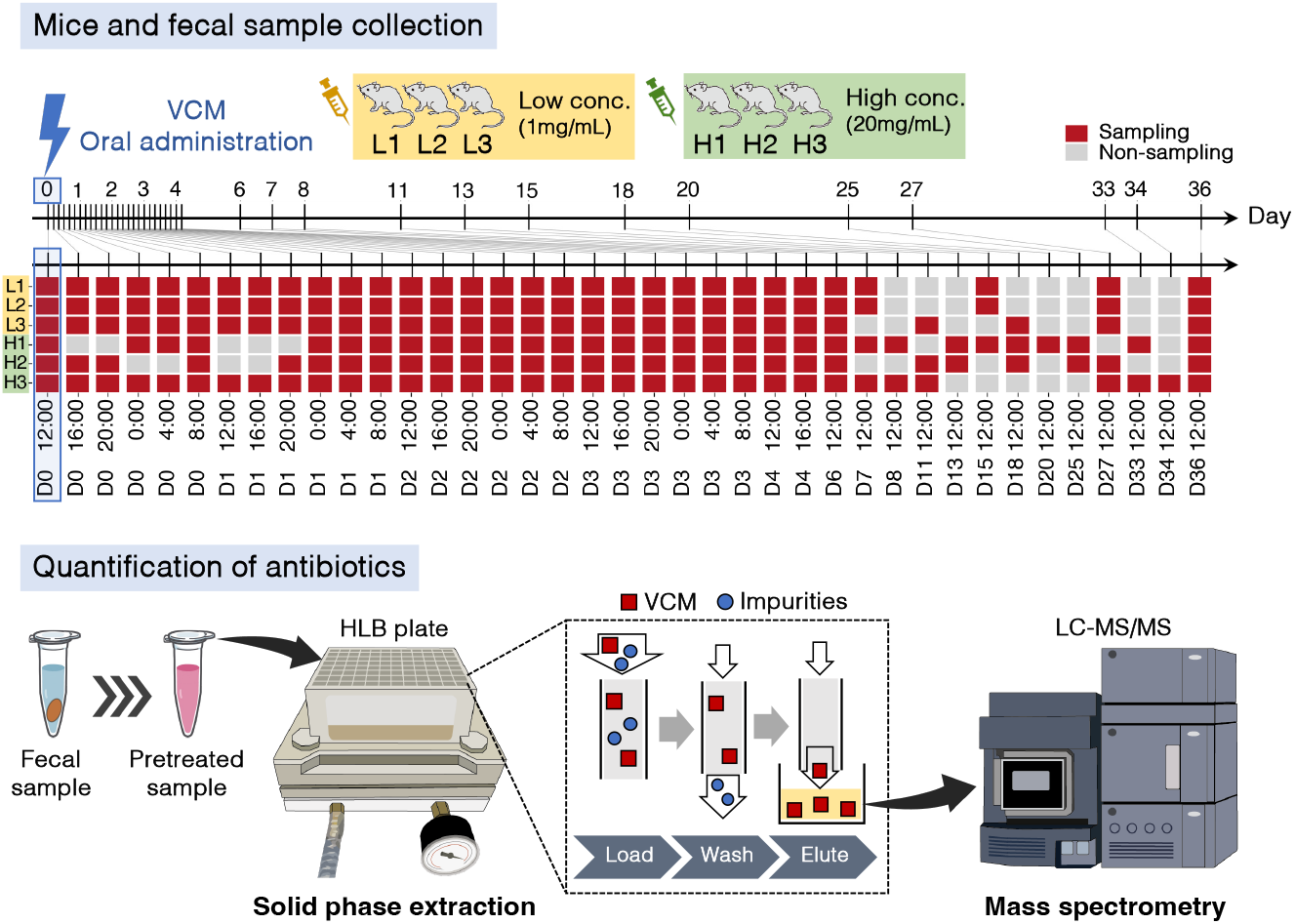
Fecal antibiotic concentration quantification protocol used in this study. C57BL/6J mice (n=3 per group) were orally administered low (1 mg/mL) and high (20 mg/mL) concentrations of vancomycin (VCM) at the initial time point (day 0, 12:00). Fecal samples were collected at a minimum interval of 4 hours (upper panel). Red cells indicate the time points at which fecal samples were collected. Antibiotic concentrations in the collected fecal samples were measured using solid phase extraction with parallel purification and mass spectrometry (LC-MS/MS) (lower panel).

### Mice and fecal sample collection

Load Wash Elute

Figure 2 shows the changes in the time course of the fecal antibiotic concentrations for each mouse. In both groups, fecal VCM concentrations gradually increased after administration, reached a peak, and then exhibited an exponential decay. The pharmacokinetic parameters derived from these data are summarized in Table 1. The area under the curve (AUC) was calculated using the trapezoidal method over the antibiotic detection period (shaded regions in Fig. 2). The elimination rate constant (*k*_*e*_) was estimated by linear regression of the logarithmic concentration from peak time (*t*_max_) to the elimination time (*t*_elim_). Compared to the low-dose group, the high-dose group consistently showed higher *C*_max_ and AUC, along with longer *t*_elim_. Since no significant differences in AUC/dose and *C*_max_/dose were observed between groups (t-test, *p >* 0.05), dose-dependent linearity in pharmacokinetics was not rejected (see Supplementary Information).

**Table 1.** Fecal pharmacokinetic parameters of antibiotics in each mouse. *C*_max_ represents the peak concentration, and AUC (area under the curve) is calculated using the trapezoidal method. *t*_app_ is the time after administration when the fecal drug concentration first exceeds the detection limit, while *t*_elim_ is the time after *t*_max_ (the time at which *C*_max_ is reached) when the fecal drug concentration first falls below the detection limit. *k*_*e*_ is the elimination rate constant estimated from measurements between *t*_max_ and *t*_elim_.

| Mouse | $C_{\max}$ [mg/g] | AUC [mg·hr/g] | $k_e$ [1/hr] | $t_{\max}$ [hr] | $t_{\text{app}}$ [hr] | $t_{\text{elim}}$ [hr] |
| --- | --- | --- | --- | --- | --- | --- |
| L1 | 1.0 | 5.6 | 0.35 | 8 | 8 | 24 |
| L2 | 0.39 | 2.9 | 0.68 | 12 | 8 | 20 |
| L3 | 0.53 | 4.4 | 0.21 | 8 | 4 | 32 |
| H1 | 14 | $3.7 \cdot 10^2$ | 0.16 | 36 | 16 | 84 |
| H2 | 11 | $2.5 \cdot 10^2$ | 0.12 | 20 | 8 | 84 |
| H3 | 3.9 | 24 | 0.11 | 8 | 0 | 52 |

**Figure 2.**
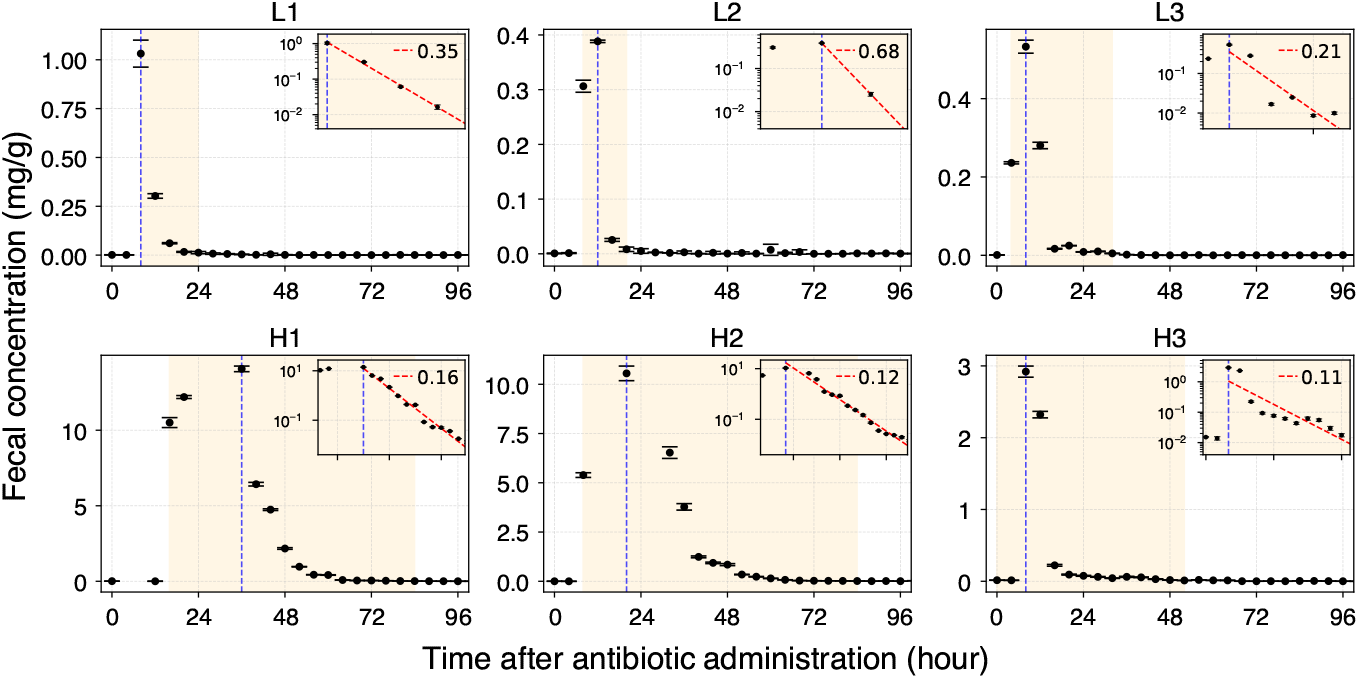
Each graph shows the time course of fecal VCM concentrations after oral administration in mice in the low group (L1–L3) and high dose group (H1–H3). Data points represent the mean and standard deviation measured by LC-MS/MS (n=3). The blue dashed line indicates the time when the maximum concentration (*C*_max_) was observed (*t*_max_). The shaded region represents the time from when the concentration first exceeded the detection limit (*t*_app_) to the time after *t*_max_ when it fell below the detection limit (*t*_elim_). The inset graph in each panel shows the log-transformed concentration changes during this period, with the red dashed line representing the linear fit result and its slope (*k*_*e*_, elimination rate constant) from *t*_max_ to *t*_elim_.

### Gastrointestinal transit model

Based on the observed pharmacokinetic characteristics, we constructed a pharmacokinetic model to describe the antibiotic transit through the gastrointestinal tract. The fecal concentration of VCM gradually increased after administration, suggesting that the antibiotic is initially retained before reaching the compartment where feces are formed. Given that mice have pouch-like structures in the stomach and cecum, we developed a mathematical model of intestinal transit, designating the stomach as the absorption compartment and the cecum as the excretion compartment (Figure 3(a)). This model assumes that VCM is instantaneously distributed in the stomach at time *t* = 0, then moves to the cecum at a constant rate, and is subsequently eliminated from the cecum following first-order kinetics. The transition processes between compartments were described using rate equations. The fecal concentration of VCM at time *t*, denoted as *C*(*t*), is expressed by the following equation (derived in the Methods section):

**Figure 3.**
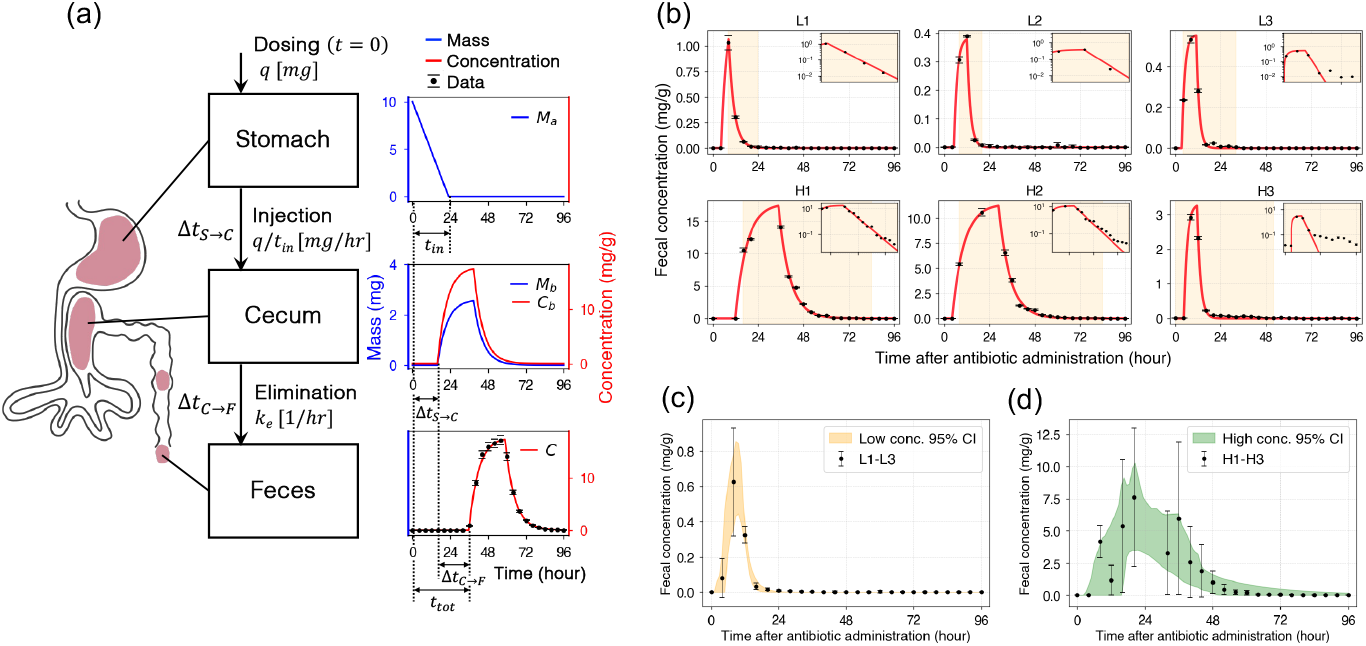
(a) Schematic representation of the intestinal transit model. The antibiotic is initially distributed in the stomach, moves to the cecum at a constant rate, and is eliminated with a rate constant *k*_*e*_. The blue and red lines indicate the mass and concentration of the antibiotic in each compartment, respectively. (b) Results of model fitting for each mouse. The red line represents the fitted model with estimated parameters (Table 2). The shaded background regions indicate the period during which the antibiotic was detected in the empirical data, and the insets show the concentration on a logarithmic scale. (c) The 95 % confidence interval of the model estimated using resampled data from the low-dose group. (d) The 95 % confidence interval of the model estimated using resampled data from the high-dose group.

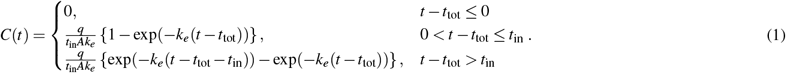

**Table 2.** Estimated parameters for each mouse. *t*_tot_ represents the total gastrointestinal transit time of the antibiotic, *t*_in_ is the duration during which the antibiotic is continuously infused into the cecum at a constant rate from the stomach, *k*_*e*_ is the elimination rate constant in the cecum, and *A* represents the mass of the cecal contents.

| Mouse | $t_{\text{tot}}$ [hr] | $t_{\text{in}}$ [hr] | $k_e$ [1/hr] | $A$ [g] |
| --- | --- | --- | --- | --- |
| L1 | 4.2 | 4.1 | 0.35 | 0.24 |
| L2 | 4.9 | 7.0 | 0.57 | 0.32 |
| L3 | 3.2 | 7.8 | 0.66 | 0.17 |
| H1 | 12 | 23 | 0.17 | 0.15 |
| H2 | 4.0 | 25 | 0.16 | 0.22 |
| H3 | 4.3 | 7.1 | 0.58 | 0.76 |

where *t*_tot_ represents the total transit time of the antibiotic through the gastrointestinal tract, *t*_in_ is the duration over which the antibiotic is infused into the cecum at a constant rate, *k*_*e*_ is the elimination rate constant of the cecum, and *A* represents the mass of cecal contents.

The fitting of the model was performed for each mouse individually, and the parameters *t*_in_, *k*_*e*_, and *A* were estimated using the *basinhopping* function in the SciPy library. The administered dose was set to *q* = 0.5 for the low-dose group and *q* = 10 for the high-dose group. Figure 3(b) shows the simulated concentration profiles using the estimated parameters for each mouse, demonstrating that the model accurately captures individual pharmacokinetic trends. The estimated parameters are summarized in Table 2.

The antibiotic transit time from the stomach to the cecum (*t*_in_) was estimated to be 4.1 to 7.8 hours in the low-dose group (L1–L3), while it was generally longer in the high-dose group (H1–H3), ranging from 7.1 to 25 hours with considerable inter-individual variability. H1 and H2 exhibited particularly long transit times of 23 to 25 hours. The elimination rate constant in the cecum (*k*_*e*_) also differed between the groups: it ranged from 0.35 to 0.66 in the low-dose group but was lower in the high-dose group (0.16–0.58), with H1 and H2 showing particularly low values (note that since the fitting was performed on the original scale, the elimination rate constants differ from those listed in Table 1, which were estimated on the logarithmic scale during the decay phase). The mass of the cecal content (*A*) was particularly high in H3 (0.76 g), but no significant differences were observed between the other individuals. Figures 3(c) and 3(d) show the results of parameter estimation using pooled data from the low and high concentration groups, respectively. Confidence intervals were constructed using 100 resampling iterations with 80 % of the data. The high-dose group exhibited greater inter-individual variability, resulting in a wider confidence interval for the model predictions.

In addition, we evaluated a first-order model in which the gastric-to-cecal transition follows first-order kinetics (Supplementary Information). The first-order model showed qualitatively similar behavior to the zero-order model, with a comparable goodness of fit to the data.

## Discussion

In this study, we captured the short time scale fecal pharmacokinetics by frequent fecal sampling and parallel antibiotic quantification. The analysis of fecal pharmacokinetics showed that in all mice, the concentration gradually increased after administration and then declined exponentially. Moreover, the fecal VCM concentration was dose dependent, consistent with findings reported in both humans^4^ and mice^11^. Based on these characteristics, we constructed the intestinal transit model with the stomach as the retention compartment and the cecum as the distribution compartment. This model represents a minimal framework that reproduces the time-delayed concentration increase and exponential decay. Fitting the model to the time series data of each mouse yielded physiologically interpretable parameters that fit with high precision.

The validity of the estimated parameters can be evaluated based on previously reported experimental data. The total gastrointestinal transit time in mice has been reported to be 6–7 hours^12^, and the mass of cecal content in C57BL/6 mice varies with diet, typically ranging from 100 to 450 mg^13^. The estimated values of *t*_tot_ and *A* for each mouse are consistent with these reports, supporting the physiological validity of the parameter estimation.

Although the ceccal elimination rate constant *k*_*e*_ and the gastric antibiotic transit time *t*_in_ are difficult to measure experimentally, the estimated values can provide insight into the motility of the gastrointestinal tract. In our model, cecal clearance (CL) represents the volume of contents expelled per unit time, which can be calculated as CL [g/hr] = *A · k*_*e*_. Similarly, gastric clearance is proportional to the inverse of *t*_in_. The calculated clearance values indicate that, except for H3, the high-dose group tended to have lower clearance than the low-dose group (see Supplementary Information). This suggests that high-dose antibiotic administration may reduce gastrointestinal motility. Comparing baseline defecation frequency and stool volume per unit time may help determine whether the observed differences in clearance are due to dosage effects or individual variation. We also examined gastric emptying under both zero-order and first-order kinetics and found that both models exhibited qualitatively similar behavior and comparable goodness of fit to the data. In humans, gastric emptying follows first-order kinetics after liquid ingestion, whereas a linear phase is observed for solid food^14^. In addition, continuous food intake tends to result in exponential gastric emptying, whereas longer intervals between meals favor zero-order emptying^15^. Thus, gastric emptying depends on the physical properties of food and feeding patterns, and in reality, a mixture of zero-order and first-order kinetics is likely involved. However, distinguishing between these mechanisms on the basis of fecal pharmacokinetics remains challenging.

This study has several limitations. First, the small sample size limited the statistical power to detect significant differences between groups. Nonlinear pharmacokinetic behaviors, such as increased total gastrointestinal transit time^16^ and decreased clearance due to antibiotic administration, need to be further investigated with additional samples. Second, since the gastrointestinal structures of mice and humans differ, adapting this model for human applications would require restructuring the compartments. In humans, the cecum is relatively small, and the colon is divided into multiple segments, which likely affects the number and roles of the necessary compartments^17,18^. Furthermore, this minimal model is applicable only to substances that are not absorbed in the gastrointestinal tract. For compounds that are metabolized or absorbed, model extensions are needed. Evaluating whether changes in fecal concentrations are consistent with fecal compartments in pharmacokinetic models explaining gastrointestinal absorption and plasma drug concentrations^19,20^ would contribute to the validation of these models. In conclusion, we characterized the detailed fecal concentration dynamics after oral VCM administration and developed a simple yet physiologically interpretable pharmacokinetic model with high accuracy. This approach not only provides insights into inter- and intra-individual variability in drug pharmacokinetics but also serves as a basis for investigating the effects of antibiotics on the gut microbiota^21,22^ and gastrointestinal motility.

## Methods

### Reagents

Vancomycin hydrochloride (VCM) was purchased from FUJIFILM Wako Pure Chemical Corporation (Biochemistry grade, Osaka, Japan). Oasis HLB 96-well plate (60 mg) used as solid phase extraction (SPE) was purchased from Waters (Milford, MA, USA). Distilled water was obtained from Millipore Milli-Q water-purification system (MSD K.K., Tokyo, Japan). Formic acid (*>* 98 %, LC/MS grade) was purchased from MSD K.K. (Tokyo, Japan). Acetonitrile (LC/MS grade) was purchased from Kanto Chemical Co., Inc. (Tokyo, Japan). An EDTA-Na2 solution (0.5 M) was purchased from Merck (Darmstadt, Germany). An EDTA extraction solution was prepared by mixing pH 4.0 citrate buffer and acetonitrile at 55:45 [v/v] and adding 0.5 M of the EDTA-Na2 solution to a final concentration of 0.2 % [v/v].

### Mice and fecal sample collection

All experiments were performed with 8-weeks old C57BL/6J male specific pathogen free mice, purchased from CLEA Japan. All mice were kept in isolated cages and darkened with plastic sheets at night in a Laboratory Animal Center, Keio University School of Medicine. Mice were provided with food (CL-2, CLEA Japan, Inc.) and water ad libitum. The mouse experiment consisted of three treatment groups of three mice each, depending on the concentration of VCM administered at 12:00 on day 0 (the start of the experiment is set as day -3). The first was the control experimental group (C1, C2 and C3), which received only water orally on day 0. The other two groups were the low (L1, L2 and L3) and high (H1, H2 and H3) concentration groups, and these mice received 0.5 mL of VCM dissolved in water at 1 and 20 mg/mL orally at 12:00 on day 0, respectively. Fecal samples were collected every 4 hours at 12:00, 16:00, 20:00, 0:00, 4:00, 8:00 on day -3 to day 3, at 12:00, 16:00 on day 4 and thereafter at 12:00 every 1 to 7 days until day 36. The samples were snap-frozen in liquid nitrogen prior to storage at *−*80 °C.

The ARRIVE guidelines were followed in conducting these experiments. The animal studies were approved by The Keio University Institutional Animal Care and Use Committee (15072), and we followed Institutional Guidelines on Animal Experimentation at Keio University.

### Quantification of antibiotics

#### Preprocessing

Experimental conditions for pretreatment and solid phase extraction (SPE) followed the study of Opris *et al*.^23^ A piece of weighed fecal sample was dipped in 0.5 mL of the EDTA extraction solution^24^, and the feces was ground with pellet pestle homogenizers in microtubes. Then, ultrasonic extraction was performed at 45 °C, 40 kHz for 10 min, followed by centrifugation at 15,000 rpm for 3 min. The supernatant was collected and diluted to 4 mL with pH 4.0 formic acid buffer.

#### Solid phase extraction

The HLB cartridge was preconditioned by passing 2 mL of methanol, followed by 2 mL of distilled water. Thereafter, 4 mL of each sample was loaded 1 mL into the cartridge and rinsed with 2 mL of pH 4.0 formic acid buffer. The cartridge was then vacuum dried for 10 minutes, and eluted with 1 mL of 60 % [v/v] methanol/water solution^25^. Finally, the elution was filtered through a 0.22 µm syringe filter (PVDF membrane, Merck).

For the determination of VCM recovery using the SPE method, the 10, 20, 30, 40, and 50 mg of feces samples of control experimental group were spiked with 2, 4, 6, 8, and 10 µg of VCM prior to extraction, respectively. The VCM recovery was 91.5–138 %.

#### Mass spectrometry

Each solution was analyzed via Liquid chromatography/tandem mass spectrometry (LC/MS-MS) using a Xevo TQD MS (Waters Corporation) coupled with an Acquity UPLC H-class (Waters Corporation). The chromatography analysis was performed using an ACQUITY UPLC HSS T3 Columnn (100 mm length × 3.0 mm i.d., 1.8 µm particle size, Waters Corporation.). The column was maintained at 40 °C, and the flow rate and injection volume were 0.3 mL/min and 5 µL, respectively. Acetonitrile (mobile phase A), distilled water (mobile phase B), and distilled water containing 1 % [v/v] formic acid (mobile phase C) were used as the mobile phase solutions. The initial compositions of mobile phases A, B, and C were 10, 89, and 1 % [v/v/v], respectively, which were maintained for 2 min after the injection. Subsequently, the composition of mobile phase A was increased to 90 % at 3 min and maintained for 5 min. Thereafter, it was decreased to 1 % and maintained for 5 min to enable for equilibration. Mobile phase C was maintained at 10 % for the gradient cycle. The analysis was performed in the positive-electrospray ionization mode. The MS ion-source parameters were as follows: source temperature, 120 °C; desolvation temperature, 500 °C ; capillary voltage, 2.0 kV ; desolvation gas flow, 1,000 L/h ; cone gas flow, 50 L/h. The data acquisition was performed in the selected-reaction monitoring (SRM) mode. A mass transition ion pair, cone voltage (CV), and collision energy (CE) for the analyte were m/z 725.00 *>* 144.10, 30 V, and 30 eV, respectively. VCM quantification was performed by using the external calibration method. The calibration standards were prepared immediately before the analysis. The calibration curve exhibited good linearity in the standard concentration range of 0.01–1.0 µg/mL, and the *r*^2^ value was over 0.995. The lowest concentration of the calibration curve was defined as a quantification limit. Three mass spectrometries were performed on each sample.

#### Calculation of fecal concentration

The concentration of antibiotics in fecal samples was calculated by dividing the measurements obtained by using mass spectrometry with LC-MS/MS by the wet weight of each fecal sample.

### Mathematical model

The intestinal transit model was designed to explain the antibiotic concentration in the feces. In the digestive activity of mice, contents are pooled in the stomach and the cecum. The rate of inflow from the stomach into the cecum and the rate of discharge from the cecum were described by a function using the antibiotic mass in each organ, and the antibiotic concentration in the feces was described by solving differential equations given initial conditions.

Let *M*_*a*_(*t*) and *M*_*b*_(*t*) be the masses of antibiotics in the stomach and cecum, respectively. Assuming that *q* [mg] of antibiotic reaches the stomach instantaneously at *t* = 0 and that the time until the stomach contents are completely emptied is constant and expressed as *t*_in_, the dynamics of the antibiotic mass in the stomach can be modeled as follows:

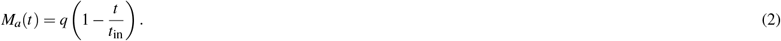

Let Δ*t*_*S→C*_ be the time from when the contents leave the stomach to when they reach the cecum through the small intestine. When Δ*t*_*S→C*_ *< t* ≤ Δ*t*_*S→C*_ + *t*_in_, the antibiotic is injected and eliminated simultaneously in the cecum, and when *t > t*_*S→C*_ + *t*_in_, only elimination occurs. Therefore, the antibiotic mass in the cecum *M*_*b*_(*t*) is as follows:

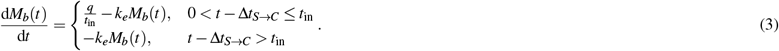

where *k*_*e*_ is the elimination rate constant. since *M*_*b*_(*t*_*S→C*_) = 0, the differential equation is solved as follows:

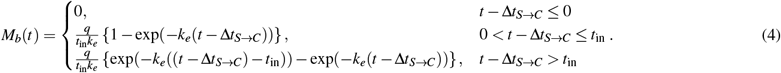

The value obtained by the experiment is the concentration of antibiotic to feces. Therefore, to consider the antibiotic concentration in the digestive tract contents, we introduce a constant *A* and consider the following antibiotic concentration *C*_*b*_(*t*) in the cecum.

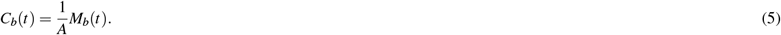

This parameter *A* can be interpreted as the mass of cecal contents. Let Δ*t*_*C→F*_ be the time between leaving the cecum and being eliminated as feces. The antibiotic concentration in feces, *C*(*t*), can be expressed in terms of the antibiotic concentration in the cecum before time Δ*t*_*C→F*_,

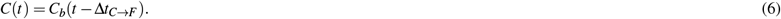

The time *t*_tot_ for the oral dose to be eliminated as feces can be expressed as *t*_tot_ = Δ*t*_*S→C*_ + Δ*t*_*C→F*_. Finally, the antibiotic concentration in feces can be calculated using the parameters *t*_tot_, *t*_in_, *k*_*e*_, and *A* as follows:

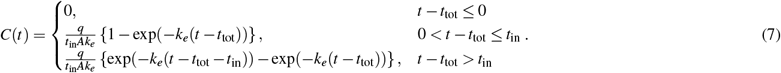

We also tested a model assuming that gastric contents are continuously diluted by gastric fluids, resulting in first-order emptying from the stomach. The equations and results are provided in the Supplementary Information.

### Parameter estimation

Parameters were estimated for each time series data for each mouse. To estimate the parameters, we minimized the following loss function, *L*(*θ*), which is given by the squared error of the data and a penalty term of errors in derivatives to control overfitting.

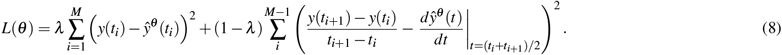

where *y*(*t*_*i*_) is the measured antibiotic concentration at sampling time *t*_*i*_ and 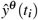 is the estimated antibiotic concentration at time *t*_*i*_ when estimated with parameter set *θ*, and *λ* is a penalty coefficient. For all estimations performed in this study, *λ* was set to 0.9. Minimization of the loss function was performed using the *basinhopping* function in the SciPy library.

## Acknowledgements

We sincerely wish to thank K.Honda and K.Atarashi for their advice on mouse experiments and A.Minowa (Keio), K. Kaida, C. Shindo, J. Noack, M. Takagi and M. Tanokura (RIKEN) for their technical support.

## Author contributions

R.M., L.T., W.S. and M.T. conceived and designed research, R.M., L.T., K.W., S.T., T.K., K.T., S.N., and W.S. performed experiments, R.M. and L.T. analyzed data, R.M., L.T., H.T., W.S. and M.T. interpreted results of experiments, R.M. prepared figures, R.M. and S.T. drafted manuscript, R.M., L.T., H.T., K.W., S.T., T.K., K.T., S.N., W.S. and M.T. edited and revised manuscript, R.M., L.T., H.T., K.W., S.T., T.K., K.T., S.N., W.S. and M.T. approved final version of manuscript.

## Data availability

All data and scripts are available on Github (https://github.com/rie-maskawa/Fecal-pharmacokinetics).

