## Supplementary Information for "High-resolution fecal pharmacokinetic modeling in mice with orally administered antibiotics"

### 1. Test for linearity of pharmacokinetics

To examine whether the pharmacokinetics follow dose linearity, a statistical test was performed. If linearity holds, $C_{max}$/dose and AUC/dose should remain constant regardless of the dose. An unpaired t-test (Welch's t-test) was used to assess the statistical significance of differences in $C_{max}$/dose and AUC/dose between the low- and high-dose groups. The results are shown in Fig. S1. For both parameters, no statistically significant differences were observed at a significance level of 5% ($p<0.05$), and thus, the null hypothesis of linear pharmacokinetics was not rejected.

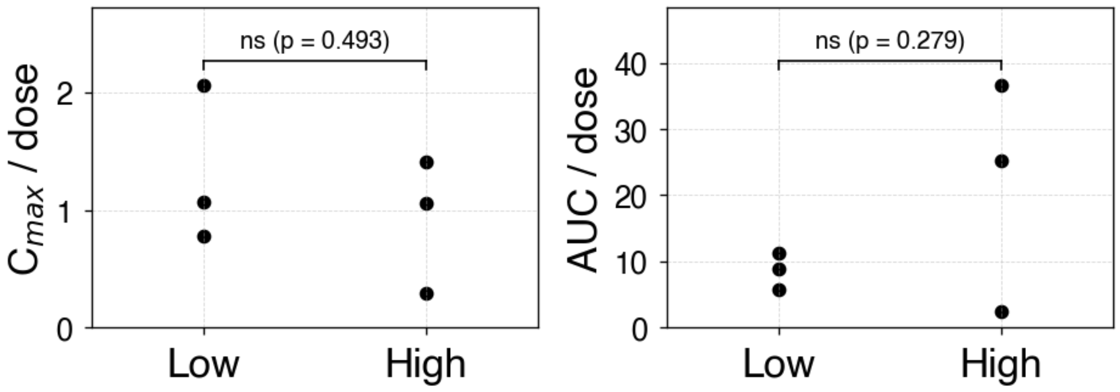


Figure S1. Results of the t-test for $C_{max}$/dose and AUC/dose

### 2. First-order model

Consider a model where gastric emptying follows a first-order rate, meaning the rate of emptying is proportional to the concentration in the stomach (Fig. S2(a)). From here on, the model where gastric emptying follows a zero-order rate will be referred to as Model 1, and the model following a first-order rate will be referred to as Model 2. In Model 2, the contents of the stomach are assumed to be continuously diluted with gastric juice and the antibiotic mass eliminated from the stomach is assumed to be exponentially decaying. If the rate constant for elimination is $k_a$, the dynamics of the antibiotic mass $M_a(t)$ in the stomach is modeled as follows:

$$\frac{dM_a(t)}{dt} = -k_a M_a(t).$$

Assuming that all of the orally administered antibiotic $q$ [mg] reach the stomach at time $t = 0$. The initial condition is $M_a(0) = q$, and the differential equation is solved as follows:

$$M_a(t) = q\exp(-k_a t).$$

The definition of the time related parameters $\Delta t_{S\to C}$, $\Delta t_{C\to F}$, $A$ and $t_{\mathrm{tot}}$ is the same as in Model 1. After $t = \Delta t_{S\to C}$, when the antibiotic reaches the cecum, injection and elimination always occur simultaneously in the cecum. Therefore, the antibiotic mass $M_b(t)$ in the cecum satisfies the following differential equation:

$$\frac{dM_b(t)}{dt} = k_a M_a(t) - k_e M_b(t).$$

where $k_b$ is the elimination rate constant from cecum. Since $M_b(\Delta t_{S\to C}) = 0$, the differential equation is solved as follows:

$$M_b(t) = \begin{cases} 0, & t \le \Delta t_{S\to C} \\ \dfrac{k_a q}{k_e - k_a}\left\{\exp\left(-k_a(t - \Delta t_{S\to C})\right) - \exp\left(-k_e(t - \Delta t_{S\to C})\right)\right\}, & t > \Delta t_{S\to C} \end{cases}.$$

In the following, $C_b(t)$ and $C(t)$ are defined as in Model 1.

$$C_b(t) = \frac{1}{A} M_b(t),$$

$$C(t) = C_b(t - \Delta t_{C\to F}).$$

As in Model 1, denoting $t_{\mathrm{tot}} = \Delta t_{S\to C} + \Delta t_{C\to F}$, the final description of the antibiotic concentration in feces in Model 2 is as follows:

$$C(t) = \begin{cases} 0, & t \le t_{\mathrm{tot}} \\ \dfrac{k_a q}{A(k_b - k_a)}\left\{\exp\left(-k_a(t - t_{\mathrm{tot}})\right) - \exp\left(-k_b(t - t_{\mathrm{tot}})\right)\right\}, & t > t_{\mathrm{tot}} \end{cases}.$$

The model fitting was performed individually for each mouse (the parameter estimation method is described in the Methods section). Figure S2(b) shows the simulated concentration profiles using the estimated parameters for each mouse. Similar to Model 1, Model 2 accurately captures individual pharmacokinetic trends. When comparing the average model fit using RMSE, Model 1 generally achieved higher accuracy, but Model 2 showed a slightly better fit for L1.

$$RMSE = \sqrt{\frac{1}{n}\sum_{i=1}^{n}\left(y(t_i) - \hat{y}(t_i)\right)^2}$$

Table S2. RMSE of Model 1 and Model 2

| | L1 | L2 | L3 | H1 | H2 | H3 | All |
|---|---|---|---|---|---|---|---|
| Model1 | $3.8\cdot10^{-3}$ | $4.6\cdot10^{-3}$ | $5.5\cdot10^{-3}$ | $5.3\cdot10^{-1}$ | $1.6\cdot10^{-1}$ | $3.2\cdot10^{-2}$ | $2.1\cdot10^{-1}$ |
| Model2 | $3.2\cdot10^{-3}$ | $8.6\cdot10^{-3}$ | $1.6\cdot10^{-2}$ | 1.5 | $5.1\cdot10^{-1}$ | $1.5\cdot10^{-1}$ | $6.0\cdot10^{-1}$ |

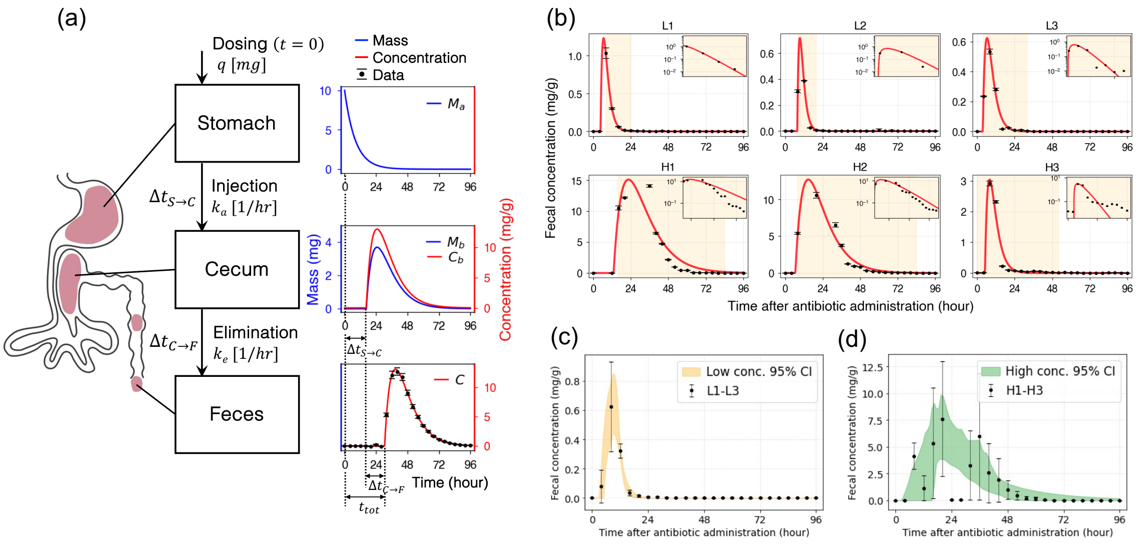


Figure S2. (a) Schematic representation of the first-order model. The antibiotic is initially distributed in the stomach, moves to the cecum with a rate constant $k_a$, and is eliminated with a rate constant $k_e$. The blue and red lines indicate the mass and concentration of the antibiotic in each compartment, respectively. (b) Model fitting results for each mouse. The red line represents the fitted model with estimated parameters (Table S1). Shaded background regions indicate the period during which the antibiotic was detected in empirical data, and the insets show the concentration on a logarithmic scale. (c) The 95 % confidence interval of the model estimated using resampled data from the low-dose group. (d) The 95 % confidence interval of the model estimated using resampled data from the high-dose group.

Table S1. Estimated parameters of the first-order model for each mouse. $t_{\mathrm{tot}}$ represents the total gastrointestinal transit time of the antibiotic, $k_a$ is the elimination rate constant in the stomach, $k_e$ is the elimination rate constant in the cecum, and $A$ represents the mass of the cecal contents.

| Mouse | $t_{tot}$ | $k_a$ | $k_e$ | $A$ |
|---|---|---|---|---|
| L1 | 4.6 | 0.41 | 0.62 | 0.12 |
| L2 | 7.7 | 0.60 | 0.60 | 0.26 |
| L3 | 3.6 | 0.37 | 0.37 | 0.29 |
| H1 | 13 | 0.11 | 0.11 | 0.24 |
| H2 | 6.5 | 0.12 | 0.12 | 0.29 |
| H3 | 5.0 | 0.34 | 0.34 | 1.2 |

### 3. Clearance

In pharmacokinetics, clearance (CL) generally refers to an organ's ability to eliminate a drug. In our model, cecal CL represents the amount of content expelled per unit time and is calculated as Cecum CL [g/hr] = $A \cdot k_e$. Similarly, gastric CL is proportional to the inverse of $t_{in}$. The calculated clearance values indicate that, except for H3, the high-dose group tended to exhibit lower clearance than the low-dose group (Fig. S3).

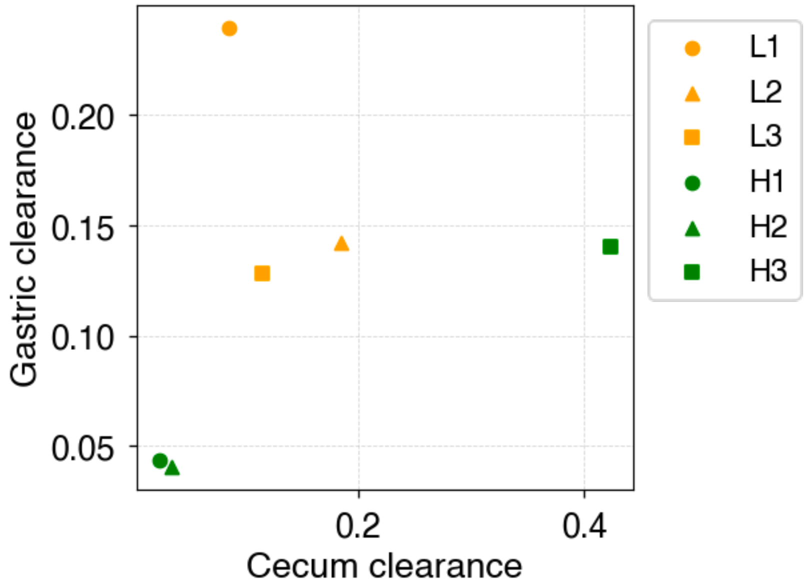


Figure S3. Cecal and gastric clearance for each mouse, calculated from the estimated parameters of the zero-order model (Model 1). Cecal clearance is computed as $A \cdot k_e$, and gastric clearance is calculated as $t_{in}{}^{-1}$.
